# Using state space models to understand trait evolution in fossil lineages

**DOI:** 10.1101/2024.07.17.603977

**Authors:** Gene Hunt, Wilmer Martinez Rivera, Melanie Hopkins, John Fricks, Beckett Sterner

## Abstract

Linear state space models provide a useful framework for investigating phenotypic evolution in fossil lineages in a wide variety of models including Brownian motion, Ornstein-Uhlenbeck processes, and models that incorporate potentially explanatory environmental covariates. A state space framework also provides access to residuals for the predicted and observed values at each time point as well as improved numerical stability. We illustrate the value of the state space approach by re-analyzing a classic dataset of diatom evolution in Yellowstone Lake. We find that number of spines is best explained by adaptation to changing solar insolation as an exogenous environmental covariate.

## 1 Introduction

The project of documenting and explaining historical patterns of evolutionary change has had enduring significance since George Gaylord Simpson introduced the ideas of evolutionary tempo and mode as a tool for bridging observation and theory across micro- and macro-evolutionary time scales (Simpson 1944). Fossil trait series provide a sequence of phenotypic measurements drawn from multiple organisms in the same lineage over a period of time, typically on the scale of tens to hundreds of thousands of years. In contrast to comparative methods, which rely on measurements of different species at particular moments, fossil trait series document historical patterns of evolutionary change in a single species over much longer durations than we can access in the present. This makes fossil trait series a uniquely valuable data source for addressing fundamental questions in evolutionary biology occurring at those scales, including why evolutionary rates of change appear to show a robust scaling with time (Harmon et al. 2021) and whether evolutionary divergence is driven by the gradual accumulation of small changes or short, rapid bursts of change that punctuate long periods of stasis (Hunt, Hopkins, and Lidgard 2015). In addition, paleobiologists have looked for signatures of other evolutionary and ecological processes, seeking to gain insights into processes of adaptation (Voje 2020; Kearns et al. 2021), extinction (Brombacher et al. 2017; Brombacher et al. 2018), stasis (Antell et al. 2021) and parallel evolution (Stuart, Travis, and Bell 2020).

Since evolution is stochastic and involves the interaction of multiple processes such as selection and drift, statistical modeling is essential to reliably estimating evolutionary rates and classifying fossil trait series into biologically significant patterns or modes of change (Hunt 2012; Gingerich 2019). In addition, the sample sizes of fossil lineage data are often small relative to the potential complexity of the system, so it is especially important to make effective use of the available information. The appropriate interpretation of evolutionary rates in a system, for example, is sensitive to the underlying mode of change: for a lineage evolving according to classical Brownian motion, we can understand its rate of change as the variance of random fluctuations it undergoes between points of time, but for a lineage exhibiting a linear directional trend in addition to Brownian motion we also need to incorporate the magnitude of that linear tendency (Hunt 2012). Numerical estimation of evolutionary rates is also sensitive to model choice, and using an inadequate model for the data can generate scaling artifacts in the magnitude of rates in relation to the absolute time duration of the trait series (Hunt 2012; Gingerich 2019; Harmon et al. 2021).

In this paper, we present some practical tools and background theory for the use of linear state space models, also called dynamic linear models, in the analysis of phenotypic time series from fossil lineages. This approach provides some key advances over previous methodological approaches. First, state space models, which apply to time series with observations at discrete time points, are grounded in continuous-time models, which are important for knowing how to handle time-varying parameters when there are uneven time steps between observations. Second, the state space framework allows for the ready evaluation of exogenous environmental covariates. Specifically, the framework provides access to residuals for the predicted and observed values at each time point, a powerful model diagnostic tool that is in standard use across other scientific disciplines. We illustrate the value of these residuals to propose a new interpretation of a classic dataset of diatom evolution in Yellowstone Lake (Edward C. Theriot et al. 2006). We also present some simulation studies emphasizing the quality of parameter estimation and model selection within this framework in comparison to previous frameworks for estimation.

We focus on univariate evolutionary models from a single species because this foundation is essential for numerically efficient and stable parameter estimation as paleontologists start to increasingly take advantage of high-throughput specimen processing, including machine-learning methods for image segmentation and trait extraction (Porto and Voje 2020; Goswami and Clavel 2024; He et al. 2024). However, the modeling framework we develop here is readily generalized to analyzing multivariate measurements from variable numbers of individual specimens at each time point. This approach also demonstrates the broad relevance of stochastic integral models to analyzing fossil lineage datasets, building on prior work by Reitan (Reitan, Schweder, and Henderiks 2012) and is complementary to stochastic integral models for species richness (Hannisdal and Liow 2018; Reitan and Liow 2019) and comparative phylogenetic datasets (Blomberg, Rathnayake, and Moreau 2020; Turley 2020).

## 2 Methods

In this section, we briefly present the definition of a linear state space model, sometimes known as a dynamic linear model. We introduce the general structure of the model along with some adaptations and quantities that this formulation facilitates. Then, we show how observing an Ornstein-Uhlenbeck (OU) process at discrete time points can be formulated as a linear state-space model. In particular, we make clear the relationship between the continuously-defined model and the discretely-sampled observations. We also discuss the relationship of other familiar models, such as an undirected random walk (RW), random walk with a directional trend (DT), and stasis (ST) to the linear state space models in the appendix.

### 2.1 Definition of a Linear State Space Model

The linear state space model consists of two recursively defined equations: the system equation and the observation equation. We will focus on the uni-dimensional case for each equation, although multivariate extension is straightforward. The system equation is a linear stochastic equation describing the underlying dynamics of a phenomena, such as the change of the level of a trait measurement for a population at a given time. The system equation can be written as

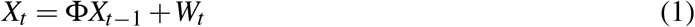

for *t* = 1, …, *T* where *W*_1_, …,*W*_*T*_ are a sequence of independent and identically distributed normal random variables with zero-mean and variance 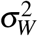. This is an autoregressive model with coefficient Φ. The model is called autoregressive because the sequence could be viewed as having the current time observation depend on the previous observation as a covariate. The observation equation is then defined as

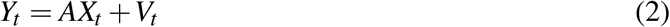

for *t* = 1, …, *T* where *V*_1_, …,*V*_*T*_ is a sequence of independent and identically distributed normal random variables with zero-mean and variance 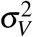 which is also independent of the sequence *W*_1_, …,*W*_*T*_. This sequence of *Y*_1_, …*Y*_*T*_ will model the observed data and is a linear transformation by *A* of the system process, *X*_*t*_, plus an additional observation error, *V*_*t*_. Note that the notation here follows the book by Shumway and Stoffer, where detailed calculations associated with this model can be found (Shumway, Stoffer, and Stoffer 2000).

In the sub-section that follows and in the appendix, we will show how a number of familiar trait evolution models fit into this framework, especially when a few features are added. This linear state space model is the framework for calculating model likelihoods using the renowned Kalman filter. The Kalman filter recursively calculates the conditional distribution of *X*_*t*_ given observations *Y*_1_, …,*Y*_*t*_; this means that if you have the Kalman filter calculated up to time *t* − 1 and receive a new observation *Y*_*t*_, then the conditional distribution of *X*_*t*_ given the data up to time *t* can be calculated without passing back through all of the previous data. A by-product of calculating the Kalman filter is that the likelihood function can be calculated with one pass through the data, so that the computational efficiency of this calculation is of order *T*. The likelihood function for a linear state space model can also be calculated using a multivariate normal distribution of size *T*, which has been the standard approach in previous paleontological research (Hunt, Bell, and Travis 2008; Voje 2020); however, that approach requires calculating the inverse of a *T ×T* matrix. Hence there can be a significant speed up and improved stability using the Kalman filter algorithm to calculate the relevant likelihood function, especially for time-series with many samples.

The Kalman filter approach also makes prediction relatively straightforward. The filter calculates the conditional mean and variance of *X*_*t*_ given *Y*_1_, …,*Y*_*t*_, giving us access to a “best guess” for *X*_*t*_ with the data up to that time. This also allows us to predict, i.e. to find the conditional mean and variance, of the next observation *Y*_*t*+1_ given the observations up to time *t* (i.e. *Y*_1_, …*Y*_*t*_), which is known as the one-step ahead predictor of the data and also allows us to construct residuals for our data after having fit it to a specific state space model. Again, using the notation of Shumway and Stoffer, the standardized residuals would be defined as

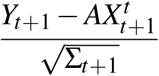

for *t* = 1, …, *T* −1, where Σ_*t*+1_ is the conditional variance of *Y*_*t*+1_ given *Y*_1_, …,*Y*_*t*_, which is calculated as part of the Kalman filter algorithm. Residuals in a time series context are an important tool for discerning the quality of fit of a model, in a manner similar to their use in regression analysis. It is a way to “approximate” the sequence *V*_*t*_, and we therefore expect them to be approximately independent and identically distributed.

In addition to facilitating likelihood function calculations and predictions of both the system equation model and future observations, the linear state space model facilitates regression with exogenous variables, which is especially important when considering the effects of environmental variables on trait mean dynamics. These can enter either through the system equation or the observation equations. In addition, the parameters defining the linear state space model can be time-varying. These modifications give

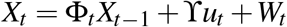

and

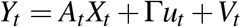

where *u*_*t*_ is an *r ×* 1 column vector of exogenous variables, typically what we think of as covariates, for each *t*, and ϒ and Γ are the coefficients that convert changes in the input variables into changes in *X*_*t*_ and *Y*_*t*_. In this context, we can still use the Kalman filter to calculate the likelihood and to create predictions and thus residuals to evaluate model performance. Visually examining the residuals is a useful complement for formal model misspecification tests for common fossil lineage models introduced in (Voje 2018).

### 2.2 Unbiased Random Walk as a Linear State Space model

Many commonly used trait evolution models can be represented in this state space framework. Perhaps the simplest case is that of an unbiased random walk, which we develop here as an example. Under this model, the trait value at one time step (*X*_*t*_) is equal to the value at the previous time step (*X*_*t*−1_), plus an evolutionary perturbation (*W*_*t*_). The autoregressive coefficient is unity (Φ = 1), and the variance of the *W*_*t*_ is usually called the step variance in paleontology. The state equation represents the true trait mean of the population at each time step. Since sample sizes are finite, however, we can never know the true means of our samples. The calculated trait means that we observe, *Y*_*t*_, reflects the true population mean (*X*_*t*_), plus sampling noise (*V*_*t*_). In most paleontological cases, our measurements of traits are noisy but unbiased, and so *A* = 1. For trait means, the observational variance of *V*_*t*_ for each sample will be the within-sample trait variance divided by the number of individuals measured in that sample.

### 2.3 Ornstein-Uhlenbeck as a Linear State Space model

The Ornstein-Uhlenbeck process is used in evolutionary studies to model adaptation (Hunt, Bell, and Travis 2008; Hansen 1997). It is often defined by its stochastic differential equation formulation.

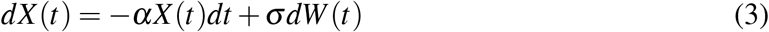

This type of differential is a short hand to define an integral equation. Such an equation is defined as a solution to sequences of difference equations:

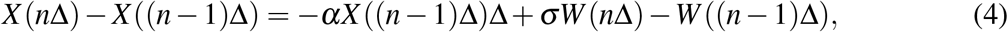

as Δ goes to zero and *n*Δ converges to some terminal time *T* and with *X* (0) defining an initial value for the equations. (This sequence is on an evenly spaced grid, which is not necessary in general as long as the grid spacing distance shrinks to zero.) The Ornstein-Uhlenbeck process has a solution (Oksendal 2013) in the form of a stochastic integral with deterministic integrand,

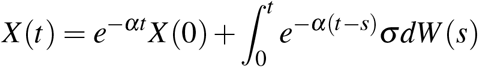

The definition of these SDEs as a limit of solutions to certain difference equations point to one possible discretization. Namely, we could re-arrange the difference equation to arrive at

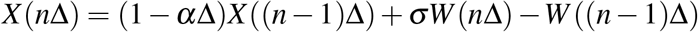

but this is only an approximate solution that accurately represents the model definition when Δ is small. However, if we take the integral solution of Equation 3 and manipulate it appropriately, we arrive at

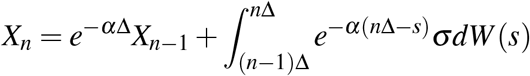

We can use Ito’s isometry to calculate the variance of the last term, which is Gaussian and has a zero mean. The variance is then

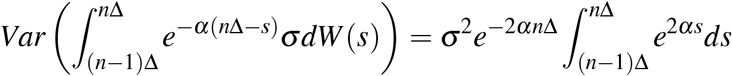

We solve the integral to arrive at the variance of the last term

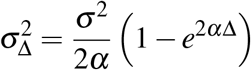

Note that when Δ is small, then a Taylor approximation will verify that this expression is approximately equal to *σ*^2^Δ corresponding to equation 4. So, we can express this discretization as

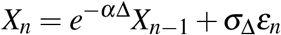

where *ε*_*n*_ is a sequence of independent standard normal random variables.

The Ornstein-Uhlenbeck model will eventually settle around zero regardless of the initial condition. We can modify this part of the model for the OU process to be centered around another constant, *θ*. The exact discrete version would be

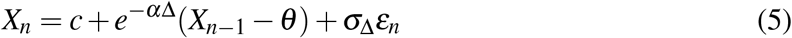

This formulation allows for one way to introduce covariates with a linear combination of covariates in place of *θ*.

### 2.4 Model estimation and selection simulations

We used simulations to validate model fitting in the state space framework using the Kalman filter compared to prior work. We also explored parameter estimation and model selection performance in the state space framework. In particular, there are two inherent observational time scales in trait series: the time step between observed samples (Δ above) and the total duration between the first and last observations. Parameter convergence may therefore depend on how increasing the sample size alters these time scales. For example, subdividing a fixed interval of time with more observations does not lead the linear trend parameter, *µ*, to converge asymptotically in the DT model. In contrast, smaller time steps are valuable for estimating the Appendix 7 provides more details on parameter estimation and model selection performance.

## 3 Application: *Stephanodiscus yellowstonensis* trait evolution

To illustrate the linear state space framework, we re-analyzed the *Stephanodiscus yellowstonensis* fossil trait series published in (Voje 2020), originally created by (Edward C. Theriot et al. 2006), using the discretized models described in Section 2.1 and implemented in the state space framework. *S. yellowstonensis* is a species of diatom endemic to Yellowstone Lake, Wyoming, United States and likely descended from the pre-existing species *S. niagarae*, which is still extant through the region (Edward C. Theriot et al. 2006). The fossil trait series is derived from 63 samples from a sediment core collected from the lake’s central basin, and it covers approximately 14,000 years ago until the present. For each sample, Theriot et al. measured 50 individuals, occasionally fewer if this number of specimens was not available. They measured three traits on each individual: valve diameter, the number of costae per valve, and the number of spines per valve. All three traits show a relatively rapid increase in values from about 12,000 to 10,000 years ago, with a slower and fluctuating decrease thereafter. As noted by (Voje 2020), all three traits are considered to be ecologically important for diatoms; valve spines, in particular, may enhance nutrient uptake and photosynthetic rate by affecting how diatoms sink through the water column. To illustrate the model-fitting methods, we focus on just one variable, spine count (Fig 1) Before analysis, spine counts were log-transformed because we consider a proportional scale to be more appropriate for the evolution of this trait. This transformation also has the effect of removing a strong correlation between the mean and variance among samples (*r* = 0.70, *P* < 1*e* − 9 for untransformed spine counts, *r* = 0.04, *P* = 0.77 after log-transformation).

**Figure 1:**
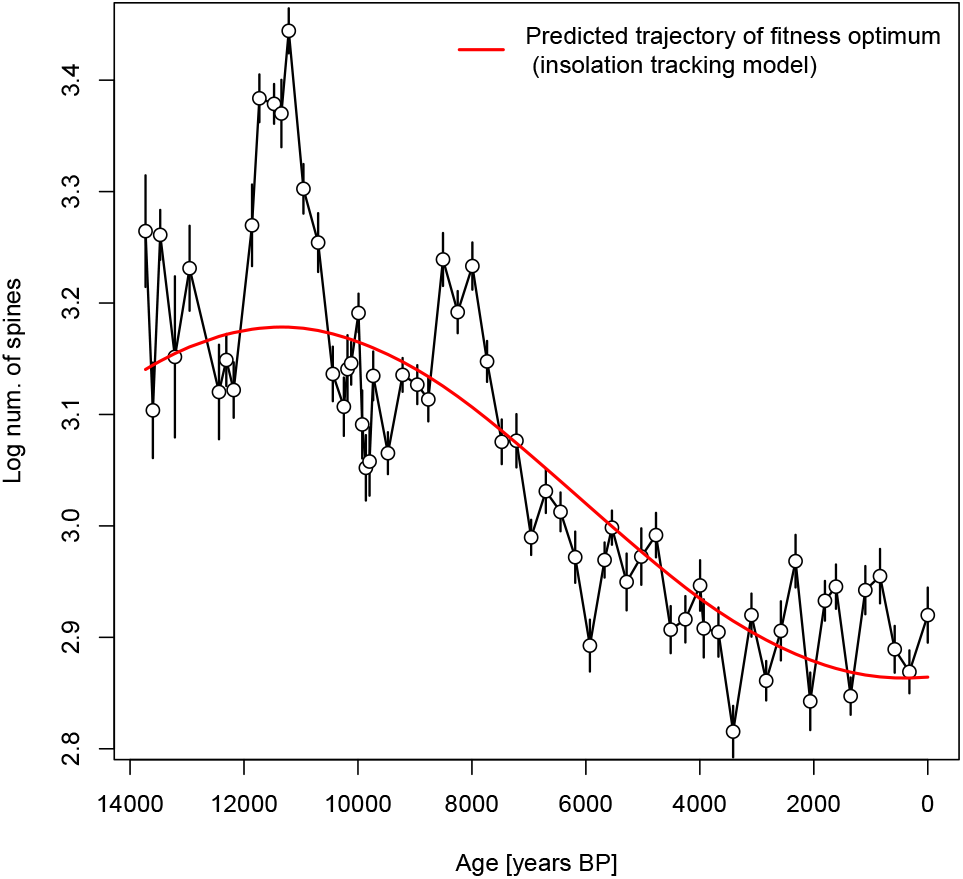
Evolutionary time-series of log spine counts for *Stephanodiscus yellowstonensis*. Open circles and vertical bars show mean values +/-one standard error on those means. The red line is the model-predicted trajectory of the fitness optimum for the best supported model (see text).

Theriot et al (Edward C. Theriot et al. 2006) posit that morphological changes in the *S. yellowstonensis* lineage track environmental changes, and these authors synthesize the available records of regional change through the study interval. Here, we quantitatively analyze two of these records. The first is the proportion of the dominant pollen type, attributable to *Pinus contortus*, as reflecting floral change in the area (Fig 2, digitized from Fig. 10 in ref (Edward C. Theriot et al. 2006)). The second environmental record we analyzed is solar insolation, which peaked around 11,000 years ago and decreased to the present day. Insolation values were taken from model output Lasker et al. (2004), using the web interface vo.imce.fr. Insolation values were computed in *W/m*^2^ at the latitude and longitude of Lake Yellowstone. As *S. yellowstonensis* blooms in summer (Edward C. Theriot et al. 2006), we used insolation values for the month of June (Fig 2).

**Figure 2:**
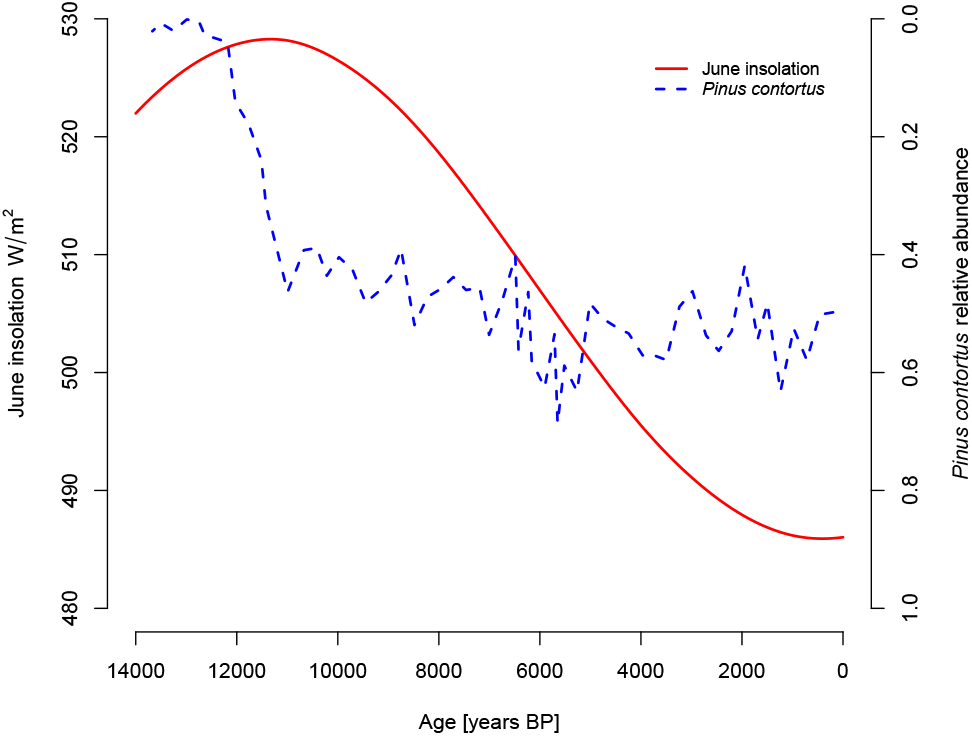
Measured environmental covariates, including June solar insolation (red solid line) and proportion of pollen attributable to *Pinus contortus* (blue dashed; note reversed axis).

The sampling times of the environmental records did not precisely match those of the trait time-series. We used linear interpolation to produce time-series of the environmental records, sampled at the same times as the traits. Both environmental records were mean-centered prior to analysis to facilitate model fitting.

Our overall model-fitting strategy started with the five models considered by Voje (Voje 2020): stasis (ST), random walk (RW), directional trend (DT), Ornstein-Uhlenbeck (OU), and decelerated evolution (DE). To test Theriot et al.’s suggestion that environmental changes influenced morphological evolution, we added OU models in which the trait optimum linearly tracks the two environmental covariates described above, June solar insolation (*OU*_*insol*_) and the proportional abundance of *Pinus contortus* pollen (*OU*_*pollen*_). Because model fits and residuals indicated a decrease in stochastic evolutionary change through the core (see Results), we considered additional models that allowed for a one-time decrease in the step variance. We specified this time to be 10,000 years ago, following Theriot et al.’s observation [(Edward C. Theriot et al. 2006), p. 45] that environmental conditions were much more stable after this date.

All the above models were fit using functions in the R package *paleoTS*, the recent update of which (v. 0.6-1) allows for fitting models via the Kalman filter and a state space model approach. Confidence intervals on parameter estimates were generated using the *dentist* package to compute approximate profile confidence intervals.

Of the five models fit by Voje, we found that DE was best supported by AICc (Table 1, models 1-5), consistent with Voje’s findings. The maximum-likelihood parameter estimate for the decay parameter of the DE model implies a roughly 7-fold decrease in the step variance over the course of the 14 kyr sequence (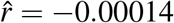, Fig. 3). Examination of residuals indicate that decrease in the stochastic component over time is an important signal in this dataset. Models without this dynamic show residuals with elevated spread early in the sequence (Fig. 4 as an example from the RW model). In contrast, residuals from the DE model are not structured in this way, showing a pattern closer to the ideal uniform spread (Fig.5). A model with a single step-down in variance at 10 ka is also consistent with these data, fitting very slightly better than the DE model (Table 1, model 6, is about 0.3 units of AICc better than DE). Parameter estimates from this model indicate that the step variance decreases from 2.77 * 10^−5^ to 1.40 * 10^−5^, about a 2-fold drop (Fig. 3).

**Table 1:** Model fits to spine counts of the Stephanodiscus yellowstonensis lineage. From left to right, columns give model abbreviations, log-likelihoods, number of parameters, AICc scores, ΔAICc scores, and Akaike weights. Model abbreviations: *RW* = random walk, *DT* = directional trend, *OU* = Ornstein-Uhlenbeck, *DE* = decelerating evolution, *RW*_*shift*_ = random walk with a shift in the step variance parameter at 10 ka, *OU*_*pollen*_ = OU model in which the trait optimum tracks the proportion of the pollen comprised of *Pinus contortus, OU*_*insol*_ = OU model in which the trait optimum tracks solar insolation, *OU*_*insol−shift*_ = OU model in which the trait optimum tracks solar insolation with a shift in the step variance at 10 ka.

| | Model | logL | K | AICc | $\Delta AICc$ | Akaike weight |
| --- | --- | --- | --- | --- | --- | --- |
| 1 | *Stasis* | 87.474 | 2 | -170.748 | 105.625 | 0 |
| 2 | *RW* | 137.003 | 2 | -269.806 | 6.567 | 0.023 |
| 3 | *DT* | 137.222 | 3 | -268.037 | 8.336 | 0.010 |
| 4 | *OU* | 138.536 | 4 | -268.383 | 7.990 | 0.012 |
| 5 | *DE* | 138.322 | 3 | -270.237 | 6.136 | 0.029 |
| 6 | *RW<sub>shift</sub>* | 138.454 | 3 | -270.500 | 5.872 | 0.033 |
| 7 | *OU<sub>pollen</sub>* | 140.355 | 5 | -269.658 | 6.714 | 0.022 |
| 8 | *OU<sub>insol</sub>* | 142.783 | 5 | -274.515 | 1.858 | 0.247 |
| 9 | *OU<sub>insol-shift</sub>* | 144.936 | 6 | -276.373 | 0 | 0.625 |

**Table 2:** Maximum-likelihood estimates (MLE) and confidence intervals (CI) for the best-fitting model: an OU process in which the position of the optimum depends linearly on the value of summer insolation and the step variance is estimated separately before 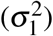 and after 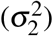 10,000 years ago. *b*0 and *b*1 are the intercept and slope of the relationship between solar insolation and the trait optimum, *α* represents the force of attraction to that optimum, and *anc* is the estimated trait value at the start of the time-series.

| | *anc* | $\sigma_1^2$ | $\sigma_2^2$ | $\alpha$ | *b0* | *b1* |
| --- | --- | --- | --- | --- | --- | --- |
| MLE | 3.24 | 4.82e-5 | 1.68e-5 | 0.00247 | 3.05 | 0.00744 |
| Lower 95% CI | 3.05 | 2.00e-5 | 9.55e-6 | 0.00025 | 2.98 | 0.00305 |
| Upper 95% CI | 3.43 | 3.05e-4 | 6.69e-5 | 0.00767 | 3.11 | 0.0141 |

**Figure 3:**
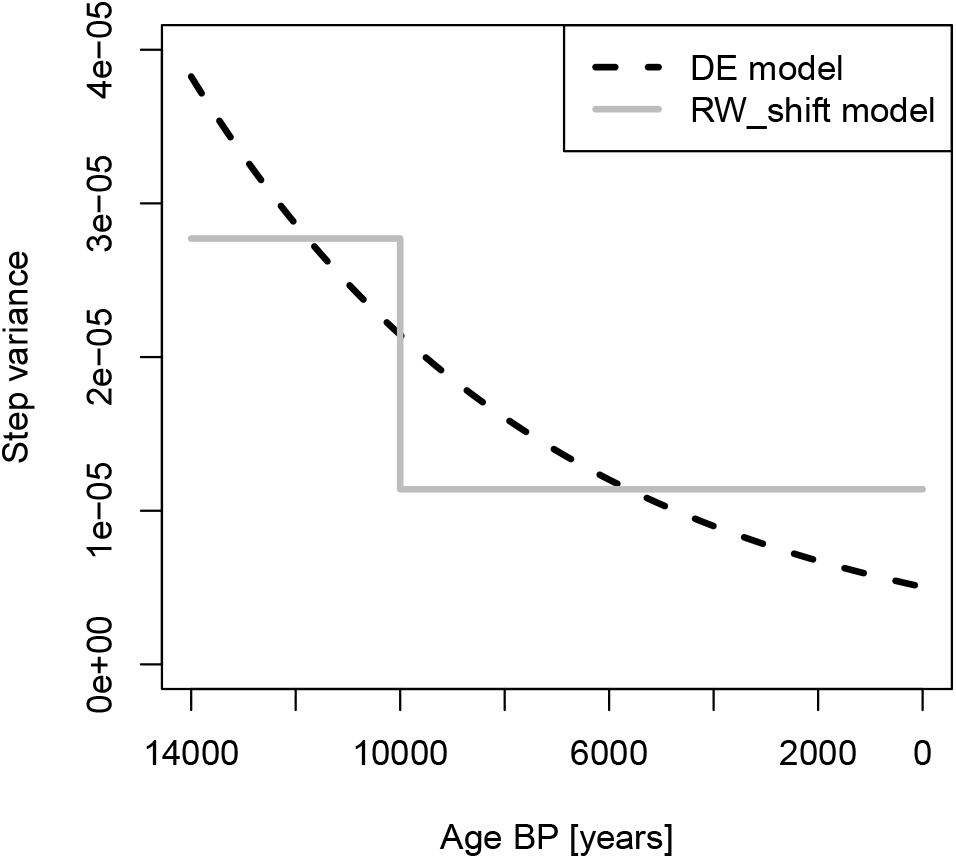
Modeled changes in the step variance predicted by the two models for which this parameter varies over time. The DE model has a step variance that exponentially decreases, whereas the RWshift model posits a single decrease in step variance that occurs at 10 Ka.

**Figure 4:**
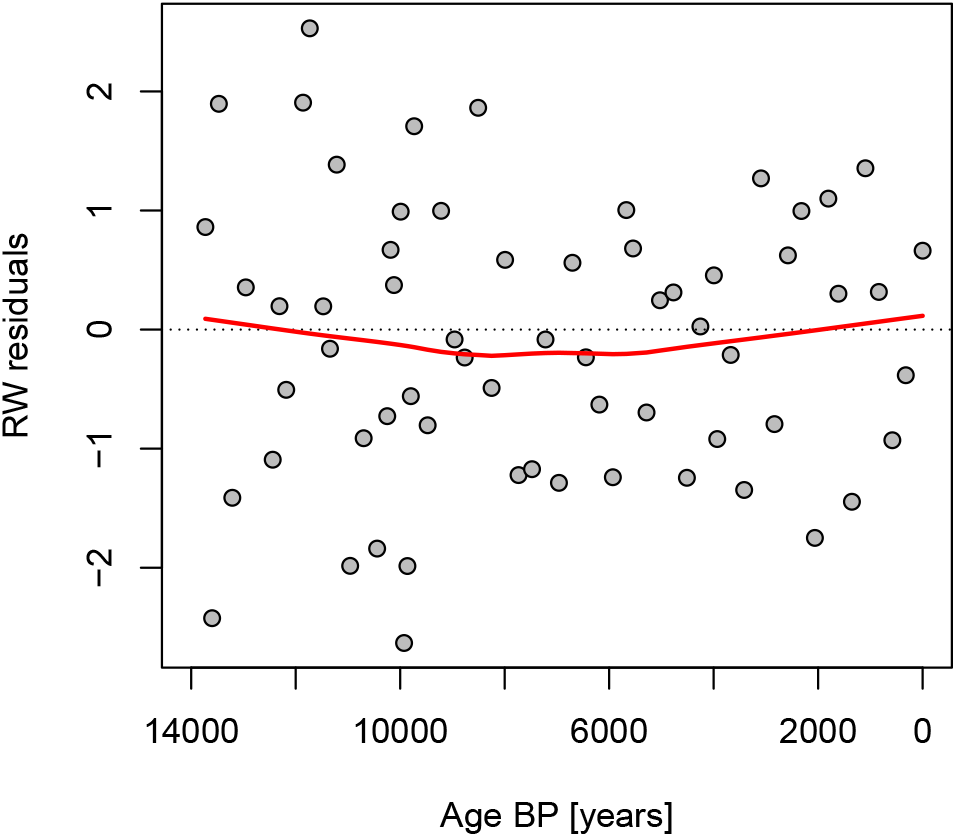
Residuals from the RWmodel. Note the greater spread of residuals early in the sequence.

**Figure 5:**
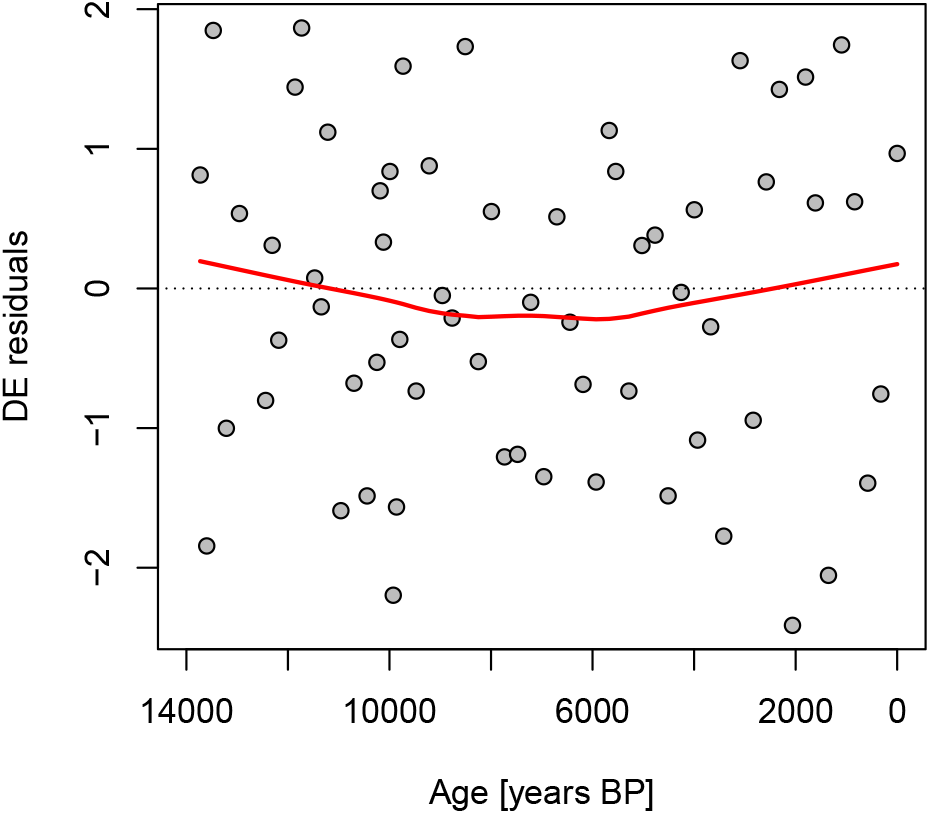
Residuals from the DE model. Note they are much less structured over time compare to those from the RW model.

Comparing the two covariate tracking models, it is more plausible that spine counts tracked June solar insolation than the pollen data (Table 1, models 8 and 7, AICc difference of 4.9). The *OU*_*inso*_ model shares the features of the OU model, except that the fitness optimum varies with solar insolation instead of being constant over time. The large increase in support between the *OU* to *OU*_*inso*_ (Table 1, ΔAICc = 6.1) is therefore a measure of the importance of solar insolation in accounting for these observations. Combining this insolation-tracking dynamic with a step decrease in stochastic variance results in a model that is best supported overall (Table 1, model 9). This model implies a dynamic where spine counts deterministically follow summer insolation, with overlaid stochastic evolution that is initially high, but then decreases later on.

## 4 Discussion

### 4.1 Using the State Space Modeling Framework

We have presented a novel framework based on stochastic integrals and linear state space models for describing, simulating, and analyzing five models for univariate trait evolution in fossil lineages. We have shown how the stochastic integral approach provides a clearer conceptual basis for relating underlying parameters stated in continuous time to models incorporating discretized sampling and observational error. In particular, we showed how a property of the sampling regime, the duration between observed time points (Δ), enters into the system equations of the Ornstein-Uhlenbeck model. Looking forward, the framework is naturally generalizable to multivariate systems.

The five base models considered here have all been implemented before for paleontological time-series, and they have been fit using maximum likelihood (**hunt 2006**; Voje 2020). The present approach, using state space models and the Kalman filter, offers an alternative means to compute these same model likelihoods. The two approaches will yield log-likelihoods that are the same (within a constant) and the resulting maximum likelihood parameter estimates are equivalent, within precision of the hill-climbing algorithm used to search for the best parameter values. Two practical benefits of using the state space approach are that the Kalman filter calculations (i) naturally produce residuals useful for assessing model adequacy, and (ii) do not require inverting a large *T* by *T* matrix, where *T* is the number of samples in the time-series. The second benefit applies mostly to rather long time-series (*T* > 100), as this inversion becomes slow and is prone to fail for very large matrices.

The analysis of spine counts in *Stephanodiscus yellowstonensis* is a good example of a typical workflow with the state space approach. An initial set of models were considered, drawn from existing theory and prior interpretations of the system under study. Model fits, as well as examination of residuals, suggest that there are two important signals in the data captured by these models: (i) a decrease over time in the stochastic component of evolutionary change, and (ii) a correlation between diatom phenotypes and summer solar insolation. The Kalman filter calculations allowed us to quickly implement a model with both of these components, which turned out to be the best supported among those considered. The modular nature of the Kalman filter can thus facilitate model development, as it is allows users to easily combine evolutionary components into new models of trait evolution.

We added a single step-down in variance, rather than the exponential decrease of the DE model, to the OU covariate-tracking model. The step decrease in variance is slightly favored over the DE model, but the decision was also a practical one; its incorporation into the OU with covariate tracking is more straightforward. Given the near-equivalance in model support between the DE and this discrete shift (Table 1), it is unlikely that these data could discriminate between the two different ways of modeling a reduction in the step variance over time.

### 4.2 Microevolution in *S. yellowstonensis*

The set of evolutionary models considered here are usually interpreted as phenomenological, not mechanistic. Phenomenological models are useful for representing, using just a few parameters, qualitatively different kinds of dynamics, such as meandering change (random walk), fluctuations around a stable mean (stasis), or directional trends. In some cases, these models can be shown to be the expected outcome of specific microevolutionary scenarios (e.g., neutral genetic drift will produce a random walk). But these models are usually interpreted descriptively, rather than as the outcome of specific microevolutionary mechanisms.

One potential exception is the OU model. Under a set of simplified but reasonable assumptions, this model describes the expected outcome of a population evolving in the vicinity of a peak in the adaptive landscape (Lande 1976). The peak corresponds to the trait value that results in highest mean population fitness. This peak is stable in the OU model, but changes with extrinsic variables in the covariate-tracking versions implemented here (*OU*_*insol*_, *OU*_*pollen*_). If the best-fitting of these *OU*_*insol*_ – were a complete description of the microevolutionary process, its parameters can be related to population genetic parameters related to the strength of natural selection (from the *α* parameter) and the effective population size (*N*_*e*_, from *σ*^2^) as described by Hunt et al. 2008 (Hunt, Bell, and Travis 2008)[page=10].

Under the above set of assumptions, the natural selection is inferred to be rather weak. Mean fitness decreases only 1% or less for population means three standard deviations away from the optimum (Table 3). Some caution should be exercised here because the timescale of adaptation is inferred to be rapid relative to temporal sampling resolution. Its half-life – the amount of time it would take the population to progress half-way to the optimum – is only about 280 years (Table 3), which is close to the median spacing between samples (258 yr). As a result, selection could be much stronger than what is estimated but we would not be able to detect it without finer temporal resolution.

**Table 3:** Estimates of microevolutionary parameters calculated from the OU model in which the trait optimum follows summer insolation, with a step down in step variance at 10 ka (model 9 in Table 1). Shown are calculations assuming low (0.1) and high (0.7) plausible values of trait heritability and 10 generations per year. *ω* is the computed variance of the population fitness function; larger values indicate broader fitness curves and therefore weaker stabilizing selection. Fitness reduction is the resulting decrease in population mean fitness between the optimal trait value and the trait values corresponding to three population standard deviations away from the optimum. Effective population size (*N*_*e*_) is computed separately before and after 10 ka.

| $h^2$ | $\omega$ | Fitness reduction | Half-life of adaptation (gen) | $N_e$ (before 10 ka) | $N_e$ (after 10 ka) |
| --- | --- | --- | --- | --- | --- |
| 0.1 | 9.37 | 1.1% | 2,802 | 482 | 1,384 |
| 0.7 | 65.7 | 0.16% | 2,802 | 3,375 | 9,686 |

The other population genetic parameter that can be calculated is the effective population size, *N*_*e*_. This parameter determines the magnitudes of change due to genetic drift in these models. Drift is more potent in smaller populations and thus lower *N*_*e*_ correspond to larger stochastic changes (= higher step variances) around the adaptive trajectory of OU models. The fit of the best model implies about a 3x increase in effective population size at 10 ka. This is consistent with a stepwise increase in the abundance of *S. yellowstonensis* observed at this time [(Edward C. Theriot et al. 2006)].

Although the direction of this change is consistent with an increase in the observed absolute abundances of this lineage, the magnitudes of estimated effective population sizes, ranging from 10^2^ to 10^4^ (Table 3), seem rather low for these unicellular algae, which can be found living at abundances high enough to produce that many individuals in just 100 liters of water, or less. (See p. 679 of (Interlandi, Kilham, and Edward C Theriot 1999)). However, it is important to note that effective population size is generally much lower than census population size, with the discrepancy between the two increasing with fluctuations in population size, differences in fitness across individuals, inbreeding, and other factors. The two literature estimates of *N*_*e*_ for diatoms are for widespread marine species and are about 10^7^, which although very high, is still orders of magnitude lower than their peak absolute abundances (Krasovec, Sanchez-Brosseau, and Piganeau 2019). It is likely that these *N*_*e*_ estimates are unrealistically low, but the population genetics of lake diatoms is not well enough investigated to be sure of this.

Assuming that the *N*_*e*_ values computed from the best model are unrealistically low, then genetic drift would not be sufficient to account for the stochastic component of spine count evolution. Therefore other factors, in addition to June insolation, will have caused changes in the position of the adaptive optimum for spine counts. Theriot et al.’s (Edward C. Theriot et al. 2006) presentation of the paleoenvironmental record provides a detailed account of environmental variation that might contribute to these insolation-independent evolutionary change. In particular, the decrease in the stochastic evolutionary component after 10 ka may be explained by the shift to more stable conditions at this time, leading to more modest selective fluctuations diatom morphology. Voje (Voje 2020) offers an alternative explanation in which ecological opportunity is initially high, perhaps because of the phenotypic changes associated with the origin of the new species *S. yellowstonensis*. With high ecological opportunity, stabilizing selection may be weakened, permitting greater variation and larger evolutionary steps. This interpretation is consistent with the observation that standing variation in spine counts is initially high and decreases steadily for the first 3 or 4 kyr of the sequence (Fig. 6). These two explanations for the reduction in stochastic evolutionary change – decreasing environmental variation and decreasing ecological opportunity – are not mutually exclusive.

**Figure 6:**
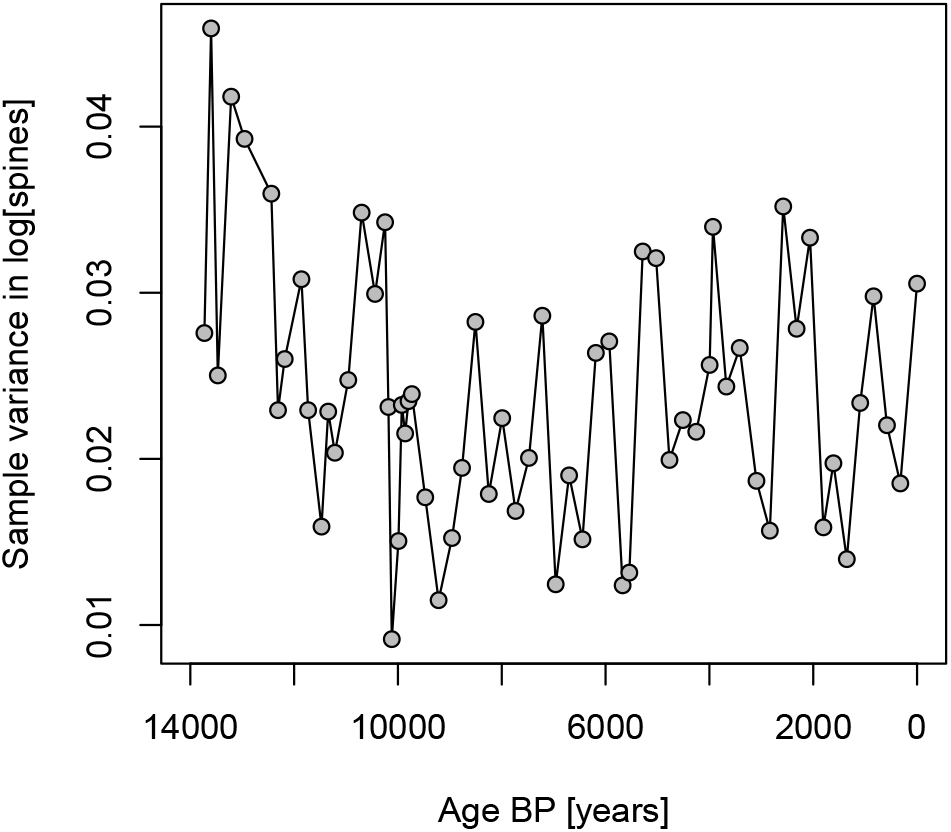
Variance in spine count decreases in the initial part of the *S. yellowstonensis* sequence.

### 4.3 Handling covariates in models of trait evolution

Although a model in which spine counts follow solar insolation as an OU process is the best supported among those considered, we caution that aspects of this model may make it less suitable for some situations. Our implementation requires an assumption that the position of the trait optimum is constant in between the time points at which we have observations. This is reasonable when, as is the case here, the covariates show point-to-point changes that are small compared to the total range of the time-series. This assumption will be less realistic for covariates that fluctuate widely on short time scales. In addition, the microevolutionary intepretation of the OU dynamics is only tenable if the sampling resolution is fine enough to potentially capture the adaptive dynamics of the population chasing the moving fitness peak. Even with the exceptional temporal resolution of the *S. yellowstonensis* data, the evolutionary dynamics may be too rapid to well-constrain the microevolutionary dynamics. In addition, this modeling approach may be prone to receiving spuriously strong support when applied to traits that show clearly directional change if analyzed with covariates that are also trended. Including the simpler model of a trend, as done here, may protect against this effect as the fewer parameters of the trend model give it an AICc advantage.

State space models are flexible enough to allow for other approaches to incorporate the effects of exogenous covariates that may be more suitable in other circumstances. For example, one could model a trait as an unbiased random walk, with an additional pulse of change that is proportional to changes in a covariate, implemented through the ϒ term of the state equation. Such a modeling approach does not attempt to capture the dynamics of a population climbing an adaptive peak and instead would be consistent with an assumption that enough time has elapsed between samples for the population to have reached the adaptive optimum. This approach would therefore would be more appropriate for trait time-series at more typical paleontological resolutions. And the use of *changes* in covariates as input variables, rather than covariates themselves, would render this approach less susceptible to trended sequences as described above.

## 5 Conclusion

The state space framework provides a practical approach for analyzing phenotypic evolution in fossil lineages that facilitates model development incorporating exogenous environmental variables based on easier access to residuals as a diagnostic tool for model fit. We highlighted some additional useful features, especially the ease of accessing time series residuals and enhanced numerical stability and efficiency. Our analysis suggested a novel biological interpretation of evolution in spine count for *S. yellowstonensis* based on stabilizing selection to changing solar insolation levels. Our focus on univariate trait models in this paper provides a foundation for expanding into more complex, multi-variate models that, for example, allow for estimation of trait covariances in a time series setting. This is essential to investigate how solar insolation that may jointly affect all three traits measured by Theriot et al.

## 6. Appendix 1

### Common models as linear state space models

#### 6.1 Stochastic Integral Models and Their Discretizations

In this section, we will discuss a number of well-known models of trait evolution in fossil lineages and give a corresponding continuous time equivalent. We will then show that each of these models can be exactly discretized to correspond to discretely observed data.

Each of these standard models may be expressed as an Ito integral with a deterministic integrand. We can then look at how each can then be expressed as a linear state space model when observed at discrete time points.

First, the definition of a Ito integral for a deterministic integrand is:

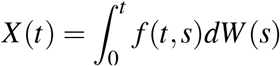

where *f* (*t, s*) is a deterministic function and *W* (*s*) is a standard Weiner process (Oksendal 2013). In other words, for each *s, W* (*s*) is a normal random variable with zero mean and variance equal to *s*. This process also has the property of independent increments, implying that *W* (*t*) −*W* (*s*) is independent *W* (*v*) −*W* (*u*) as long as (*s, t*] and (*u, v*] do not overlap. The fact that *W* (*s*) has zero mean for each *s* implies that *X* (*t*) does also. Each of the models that we examine in this manuscript is a Gaussian process and using this integral representation allows us to express the models in a unified way.

An important property of this integral, especially when calculating variances, is the Ito isometry (Oksendal 2013).

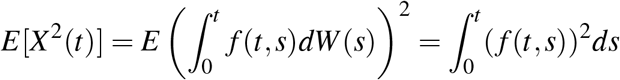

Now, we write some basic models in the form of such an integral along with their exact discretizations and approximate discretizations where appropriate.

##### 6.1.1 Random Walk

For the RW model, the deterministic function is simply a constant, *σ*, and

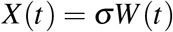

Note then, that a discretely observed version of this model, assuming equal spacing in time, would be

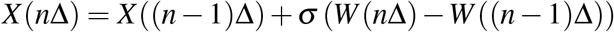

which we could we write as follows by using the fact of independent increments of Brownian motion:

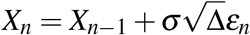

where *ε*_*n*_ are a sequence of independent standard normal random variable and Δ is the amount of time between observations. This can be derived directly from the integral definition above with the 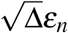 corresponding to *W* (*n*Δ) −*W* ((*n* − 1)Δ). In this discrete version, *X*_*n*_ corresponds to *X* (*n*Δ).

##### 6.1.2 Directed Random Walk

For this model, we add a deterministic linear directionality to the Random Walk model.

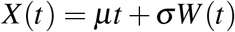

In a similar way then, a discretely observed version of this model, again assuming equal spacing in time, would be

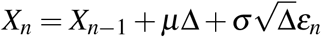

with similar interpretations as for the Random Walk.

##### 6.1.3 Decelerated Evolution

Voje introduced a model for the evolution of a trait where the step variance of a random walk declines exponentially. In other words, a model that could be describe with the recursion

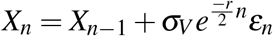

where *σ*_*V*_ and *r* are positive parameters and *ε*_*n*_ are a sequence of standard normal random variables. A natural way to write this as a stochastic integral would be

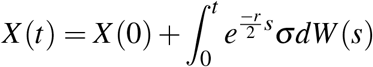

An Euler approximation of this integral would then be defined by the recursion

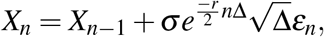

and if we identify 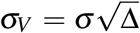 we notice that this approximation corresponds to Voje’s original definition. However, we can write down an exact discretization of the stochastic integral.

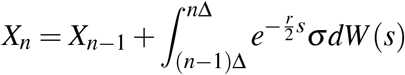

We can again look at the second term on the right and calculate the variance using Ito’s isometry

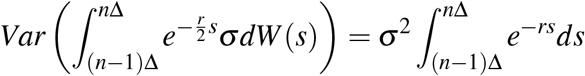

So, solving the integral on the right hand side of the above equation, we find

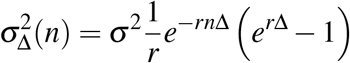

Applying a Taylor expansion with Δ small, we see that

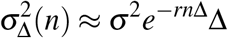

and this corresponds to the Euler approximation above.

##### 6.1.4 Stasis

The stasis model assumes that each observation is independent and identically distributed, typically with a normal distribution. So, *X*_*n*_ is normal with mean *c* and variance 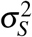, which we could write as

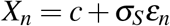

where the *ε*_*n*_ are a sequence of independent and identically distributed random variables. Effectively, there is no continuous version of this model. One way to think of this model, however, is as a discretely observed Ornstein-Uhlenbeck process with sufficient spacing between the observations relative to the parameter *α*. The Ornstein-Uhlenbeck process is stationary, meaning that after a sufficient time, the initial condition is trivially relevant and the relationship between observations at two time depends only on the distance between those observations. Since the Ornstein-Uhlenbeck process is Gaussian, the variance and covariances define the process. The covariance between two observations of a (zero-mean) Ornstein-Uhlenbeck process is given by

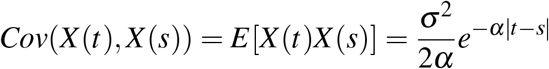

So, if the observations are sufficiently spaced this covariance is effectively zero (as long as *α* is not too small). So, the stasis model could be viewed as an Ornstein-Uhlenbeck process that is approximately stationary and sufficiently spaced with the relationship between the variances being 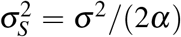 where *σ*^2^ is the infinitesimal variance of the Ornstein-Uhlenbeck model. If we look at equation 5, with *α*Δ being large, then the we see that the OU model is effectively the stasis model.

## 7 Appendix

### Simulations and Model Selection

To ensure comparability of results, we re-used code from previous studies to simulate data from the five models, including the paleoTS R package maintained by Hunt and supplementary materials from (Voje 2020). We used paleoTS to simulate data for the ST, RW, DT, and OU models and code from (Voje 2020) to simulate data for the DE model. However, we found that prior simulation studies explored parameter values and data sampling regimes in a way that varied both observational and biological time scales simultaneously (Hunt 2006; Hunt 2008; Voje 2020), making their separate effects on performance difficult to disentangle. We chose simulation parameters to feature substantial levels of model uncertainty in order to illustrate dependence on sampling scales.

For model selection, we calculated model goodness-of-fit using the corrected Akaike Information Criterion (AICc), which is modified for better performance in small sample sizes (Hunt 2008). In general, the AIC provides an unbiased estimator of a model’s expected likelihood, and picking the model with the best (i.e. lowest) AIC score will asymptotically converge on the true distribution when it is unique to a single model (Burnham and Anderson 2002). We also calculated Akaike weights for each model using the AICc scores (Wagenmakers and Farrell 2004), which approximate the probability that a model is the best out of the candidates considered. For model estimation, we used the linear Kalman filter in the state space models and compared the results of our procedures to those in PaleoTS and Voje’s code.

#### 7.1 Parameter estimation

We find that estimation accuracy and uncertainty are not uniformly influenced by sampling time scales. We show simulation results for the RW and DT models (Fig. 7), OU model (Fig. 8), and DE model (Fig. 9). We explored three scenarios for modifying sample time scales: first, increasing sampling with constant total duration (shrinking Δ); second, increasing duration and increasing sample size (uniform Δ); and third, increasing duration while holding sample size constant (increasing Δ). The ST model is not shown but is effectively an independent, identically distributed (i.i.d.) process with time-indexed observations, and so estimation of 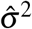 will depend on sample size but not total duration or Δ.

**Figure 7:**
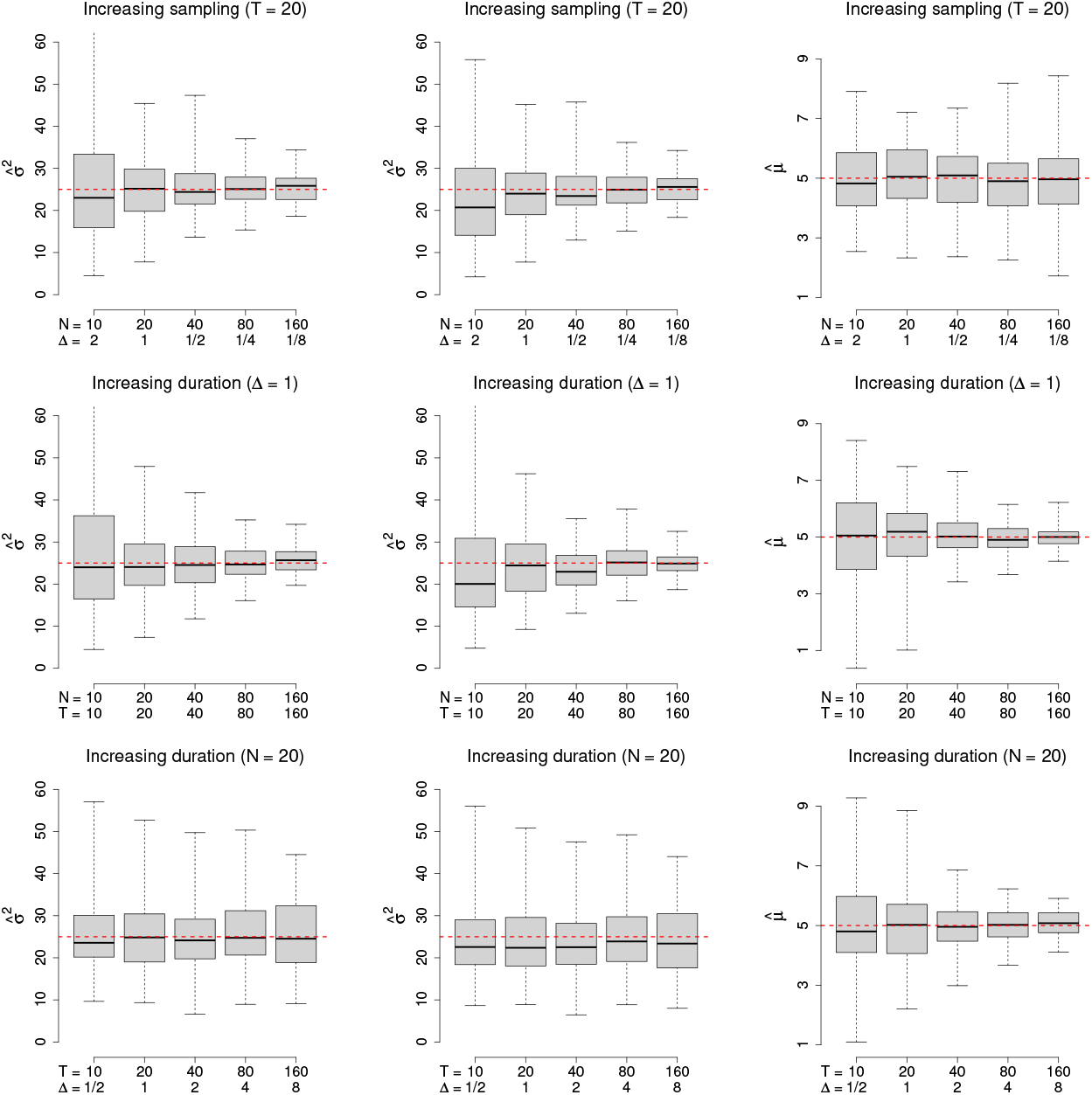
Parameter estimation for random walk and directional trend models using varying ratios of stepwise and total observational time scales. The left column shows estimation of the diffusion parameter for the RW model. The middle and right columns show estimation of the diffusion and the drift parameters, respectively, for the DT model. The true parameter values are *σ*^2^ = 25 and *µ* = 5. We use different values for the sample size *N* = {20, 40, 80, 160}, the size of the increments Δ = {1*/*8, 1*/*4, 1*/*2, 1}, and the terminal time *T* = {2.5, 5, 10, 20}. Box plots show 100 replicates

**Figure 8:**
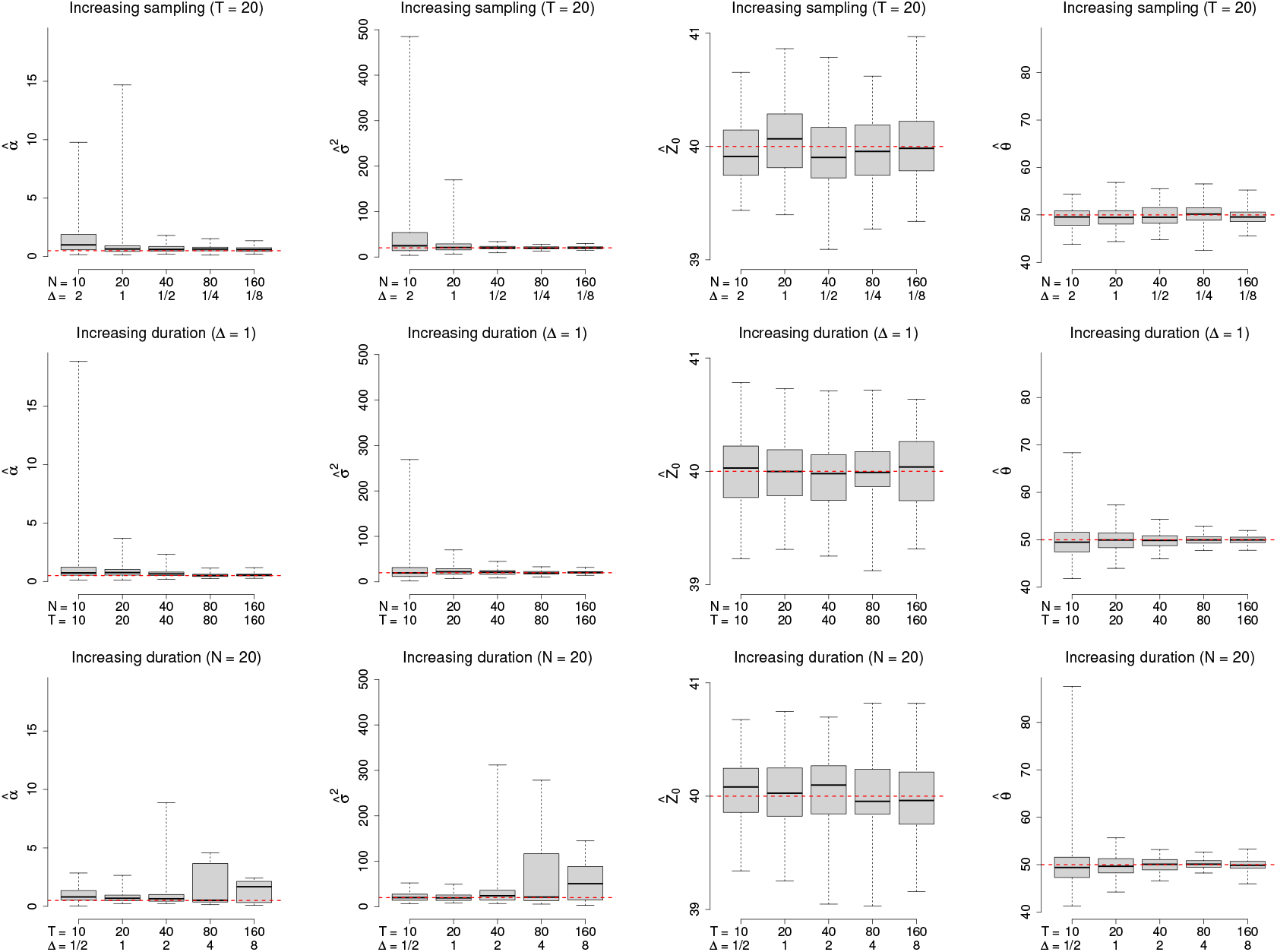
Parameter estimation on simulated data from the Ornstein-Uhlenbeck model for different combinations of sample size, time step, and total duration. Each of the columns shows parameter estimation for *α, *σ*, Z*_0_, and *θ*, respectively. True values are *α* =0.50, *σ*^2^ = 20, *Z*_0_ = 40, and *θ* = 50, as represented with dashed red lines. We varied the sample size {*N* = 20, 40, 80, 160 }, the size of the increments {Δ = 1*/*8, 1*/*4, 1*/*2, 1}, and the terminal time *T* = {2.5, 5, 10, 20}. Box plots show 100 replicates

**Figure 9:**
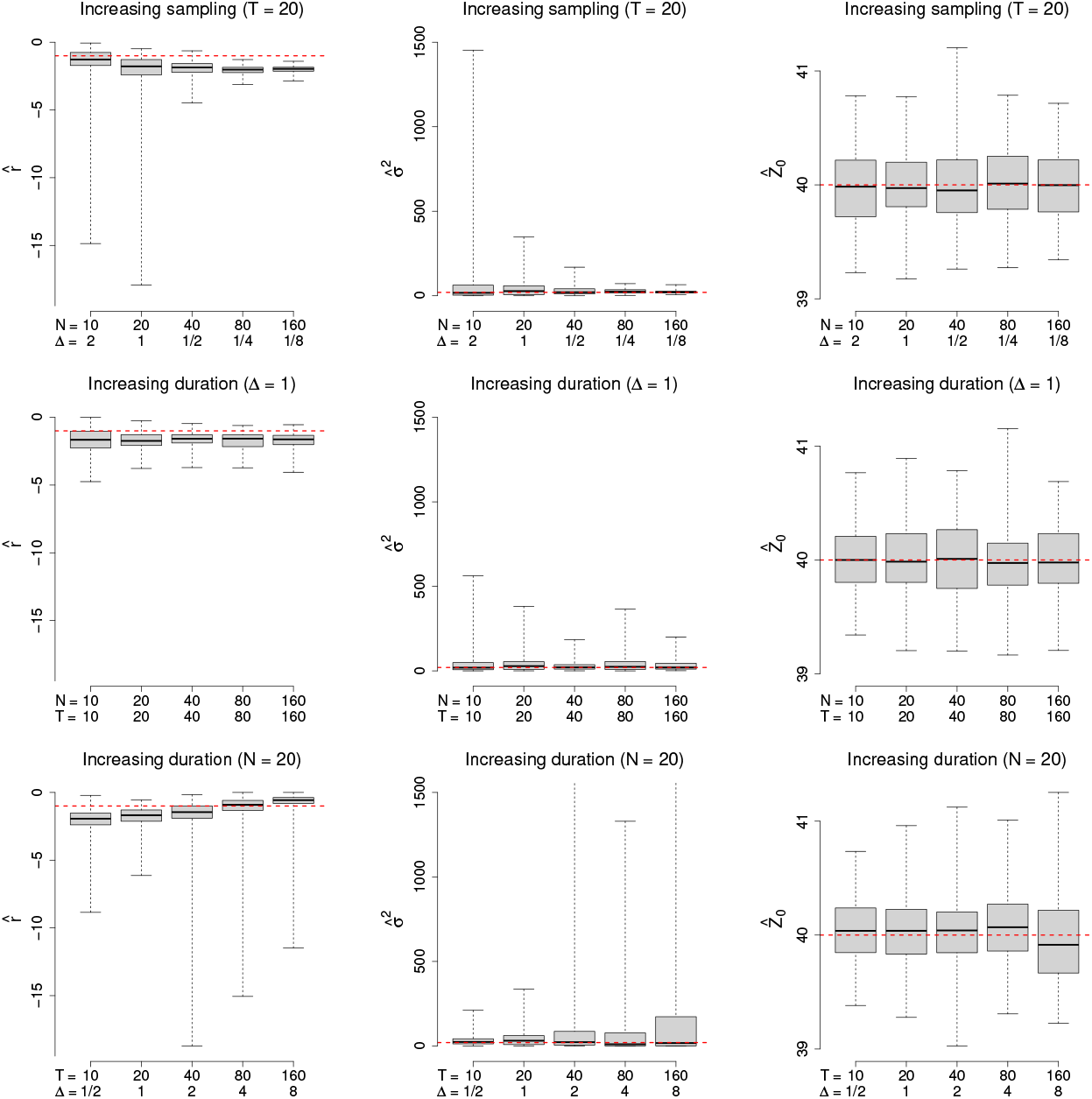
Parameter estimation for Decelerated Evolution model on simulated data using varying combinations of sample size, time step, and total duration. The columns show parameter estimation of *r, σ* and *Z*_0_, respectively. Dashed red lines show the true values of *r* =-1, 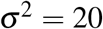, and *Z*_0_ = 40. We used sample sizes *N* = {20, 40, 80, 160}, time increments of Δ = {1*/*8, 1*/*4, 1*/*2, 1}, and terminal times *T* = {2.5, 5, 10, 20}. Box plots show 100 replicates.

**Figure 10:**
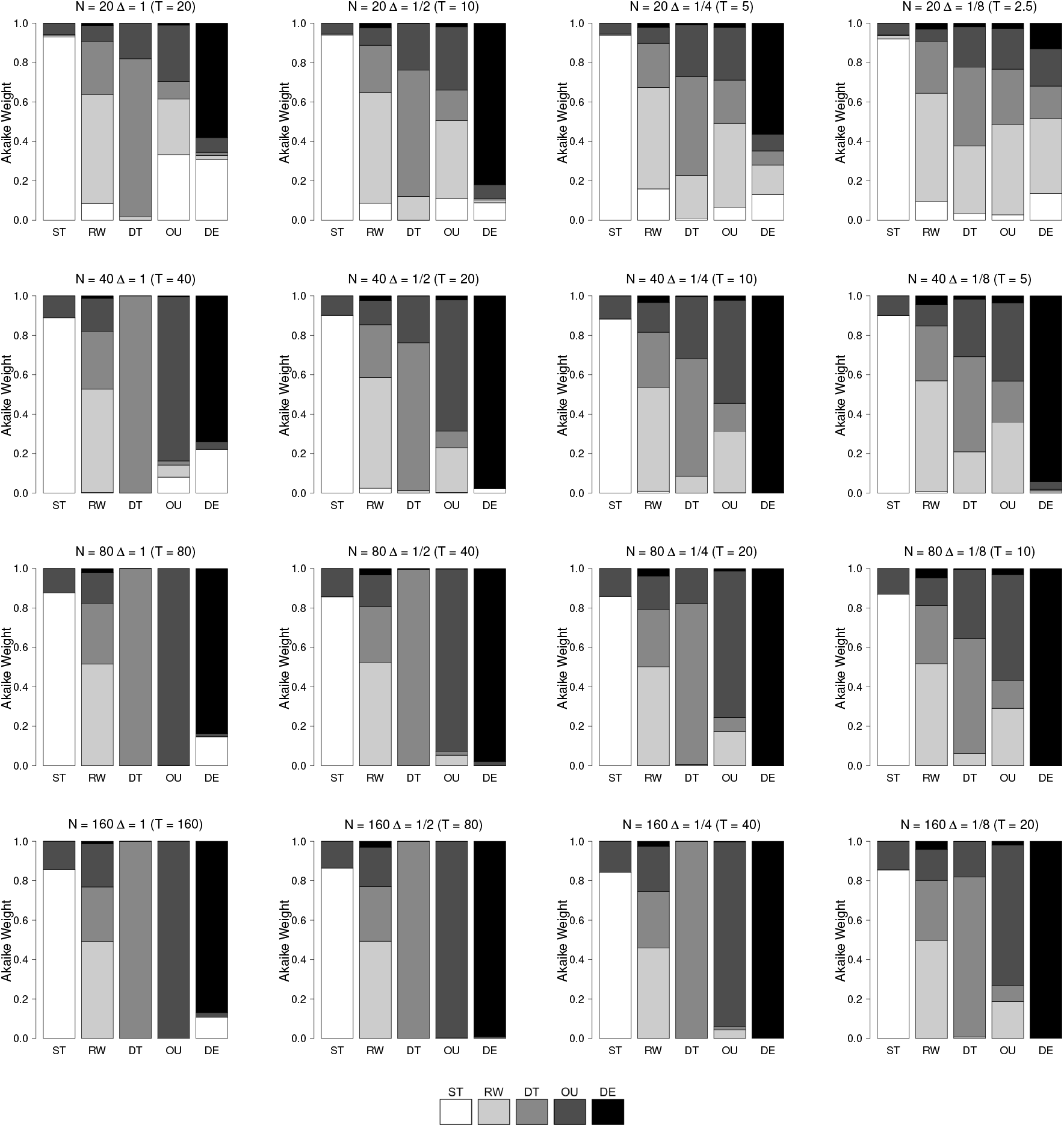
Model selection performance using the AIC criterion on simulated data for varying ratios of stepwise and total observational time scales. In each panel, the true model is labeled on the x-axis, and the stacked histogram shows the average Akaike weight for each model. Perfect model performance would show each bar as completely filled by the corresponding true model’s shade on the legend (e.g. ST as white, RW as light gray, DT as medium gray, etc.). The true model parameters are ST: *θ* = 50, *ω* = 20; RW: 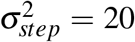 DT: *Dri f t* = 5, 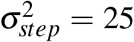 OU: *θ* = 50, *α* = 20, 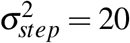 and DE: *r* = −1, 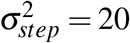. The initial condition for all the models is *Z*_0_ = 40, the variance of the evolutionary step is *V*_*p*_ = 5, and the vector of population sample size is *m* = 50. We varied sample size *N* = {20, 40, 80, 160}, the size of the increments Δ = {1*/*8, 1*/*4, 1*/*2, 1}, and the terminal time *T* = {2.5, 5, 10, 20}.

For the RW and DT models, the results in Fig. 7 show that increasing total time while keeping the sample size fixed doesn’t affect 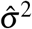 but does improve 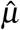. Note that the sufficient statistic for 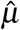 is the value of the process at the terminal point of the time series, so intermediate values don’t matter for estimation, only the end point. Increasing duration with constant Δ improves both 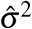 and 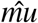, but for different reasons: 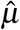 is improving because the total time observed is increasing, but 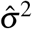 is improving because there are more steps observed.

In contrast, the OU model parameters in Fig. 8 show several different types of response to time scales. Both 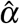 and 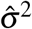 improve for both the scenarios with increasing sampling with constant duration or increasing duration. As Δ becomes larger in row 3, however, estimation gets worse because exp(−Δ) goes to zero and the process starts to look i.i.d., so that 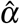 and 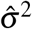 are both being fit to a normal distribution with mean *θ* and variance *σ*^2^ */*(2*α*). The initial value parameter, 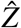, is unaffected in all three scenarios because better estimation of the restoring force, *α* can only improve precision for the initial condition up to a point. Similar to 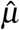 ‘s behavior in the RW and DT models, the OU equilibrium value parameter, 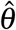, does not converge under the increased sampling intensity.

Neighboring observations are positively correlated, so adding more time points within a fixed interval gives diminishing returns for estimating the mean, but if *T* is increasing, the observations are spaced further apart and so are more independent.

For the DE model, the variance decay parameter 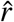 in Fig. 9 shows improved precision in the increasing sampling intensity and increasing duration, constant sampling scenarios but remains biased below the true value for the simulation setups we examined. The DE model shows phenomenologically the same behavior in 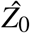 and 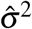 as the OU model but for different reasons. As the step variance of the process decays exponentially to zero with time, observing the process over a longer duration provides progressively less information.

#### 7.2 Model selection performance

Figure 10 shows how model selection performance, measured in terms of the average Akaike weight of the true model, varies with sampling. Columns in the figure show increasing total duration. Rows show denser sampling as total duration shrinks. Diagonals from top-left to bottom-right show increasingly dense sampling within a fixed total duration.

ST is almost exclusively conflated with OU. Increased data appears to slightly worsen false positives for OU when stasis is true, likely because the AICc has a bias for nested models toward the model with more parameters. RW is most frequently confused with DT and OU models. The average Akaike weight does not vary significantly with sampling time scales, again likely because of the AICc’s bias toward complex models. As expected, the evidence for DT improves significantly as total duration grows. OU shows improvement with greater duration and sampling density. The same is true for DE, which is mainly competitive with ST for small sample sizes.

